# RNF168-mediated localization of BARD1 recruits the BRCA1-PALB2 complex to DNA damage

**DOI:** 10.1101/2021.06.25.449923

**Authors:** John J. Krais, Yifan Wang, Pooja Patel, Jayati Basu, Andrea J. Bernhardy, Neil Johnson

**Affiliations:** Molecular Therapeutics Program, Fox Chase Cancer Center, Philadelphia, PA 19111, USA; Blood Cell Development and Function Program, Fox Chase Cancer Center, Philadelphia, PA 19111, USA

## Abstract

DNA damage prompts a diverse range of alterations to the chromatin landscape. The RNF168 E3 ubiquitin ligase catalyzes the mono-ubiquitination of histone H2A at lysine (K)13/15 (mUb-H2A), forming a binding module for DNA repair proteins. BRCA1 promotes homologous recombination (HR), in part, through its interaction with PALB2, and the formation of a larger BRCA1-PALB2-BRCA2-RAD51 (BRCA1-P) complex. The mechanism by which BRCA1-P is recruited to chromatin surrounding DNA breaks is unclear. In this study, we reveal that an RNF168-governed signaling pathway is responsible for localizing the BRCA1-P complex to DNA damage. Using mice harboring a *Brca1^CC^* (coiled coil) mutation that blocks the Brca1-Palb2 interaction, we uncovered an epistatic relationship between *Rnf168^-^* and *Brca1^CC^* alleles, which disrupted development, and reduced the efficiency of Palb2-Rad51 localization. Mechanistically, we show that RNF168-generated mUb-H2A recruits BARD1 through a BRCT domain ubiquitin-dependent recruitment motif (BUDR). Subsequently, BARD1-BRCA1 accumulate PALB2-RAD51 at DNA breaks via the CC domain-mediated BRCA1-PALB2 interaction. Together, these findings establish a series of molecular interactions that connect the DNA damage signaling and HR repair machinery.

## Main text

BRCA1’s role in maintaining genome integrity requires numerous protein interactions. The BRCA1 Really Interesting New Gene (RING), Coiled-Coil (CC), and BRCA1 C-Terminal (BRCT) domains each perform specialized functions within homologous recombination (HR) DNA repair. Significantly, germline mutations found in patients with breast and ovarian cancer often cause disruption to these domains^1,2^.

BRCA1 and BARD1 form a heterodimer with E3 ubiquitin ligase activity via their respective RING domains. BRCA1 and PALB2 each contain CC regions that directly interact with one another, permitting the formation of a larger BRCA1-PALB2-BRCA2-RAD51 (BRCA1-P) macro-complex that promotes the accumulation of RAD51 at sites of DNA damage, and consequently RAD51 filament formation^3–5^. While the BRCA1-PALB2 interaction is not necessary for the focal accumulation of BRCA1, it is required for efficient PALB2 and RAD51 foci formation, indicating that BRCA1 recruits PALB2-RAD51 to double stranded DNA break (DSB) sites. The BRCA1 BRCT domain-mediated interaction with ABRAXAS and RAP80 allows for the BRCA1 CC domain to simultaneously bind PALB2. Thus, it has been proposed that the BRCA1-ABRAXAS-RAP80 (BRCA1-A) complex is responsible for localizing PALB2 to DSBs^6^. However, several reports have demonstrated that the BRCA1-RAP80 association reduces cellular HR activity^7,8^. Moreover, RAP80 depletion was shown to have little effect on RAD51 foci formation^7^, making the role of RAP80-ABRAXAS in recruiting BRCA1-P ambiguous. Currently, the precise molecular mechanism responsible for the localization of the BRCA1-P complex has not been defined.

In response to DSBs, the E3 ubiquitin ligase RNF168 mono-ubiquitinates H2A at K13/15 (mUb-H2A), forming a binding module for several proteins, including 53BP1^9–11^. RNF8 and RNF168 can also generate K63-linked ubiquitin polymers that promote the focal accumulation of the BRCA1-A complex^12–14^. In this study, we aimed to further decipher the role of RNF168 in recruiting BRCA1-P to DSBs (**Fig. 1A**).

**Figure 1.**
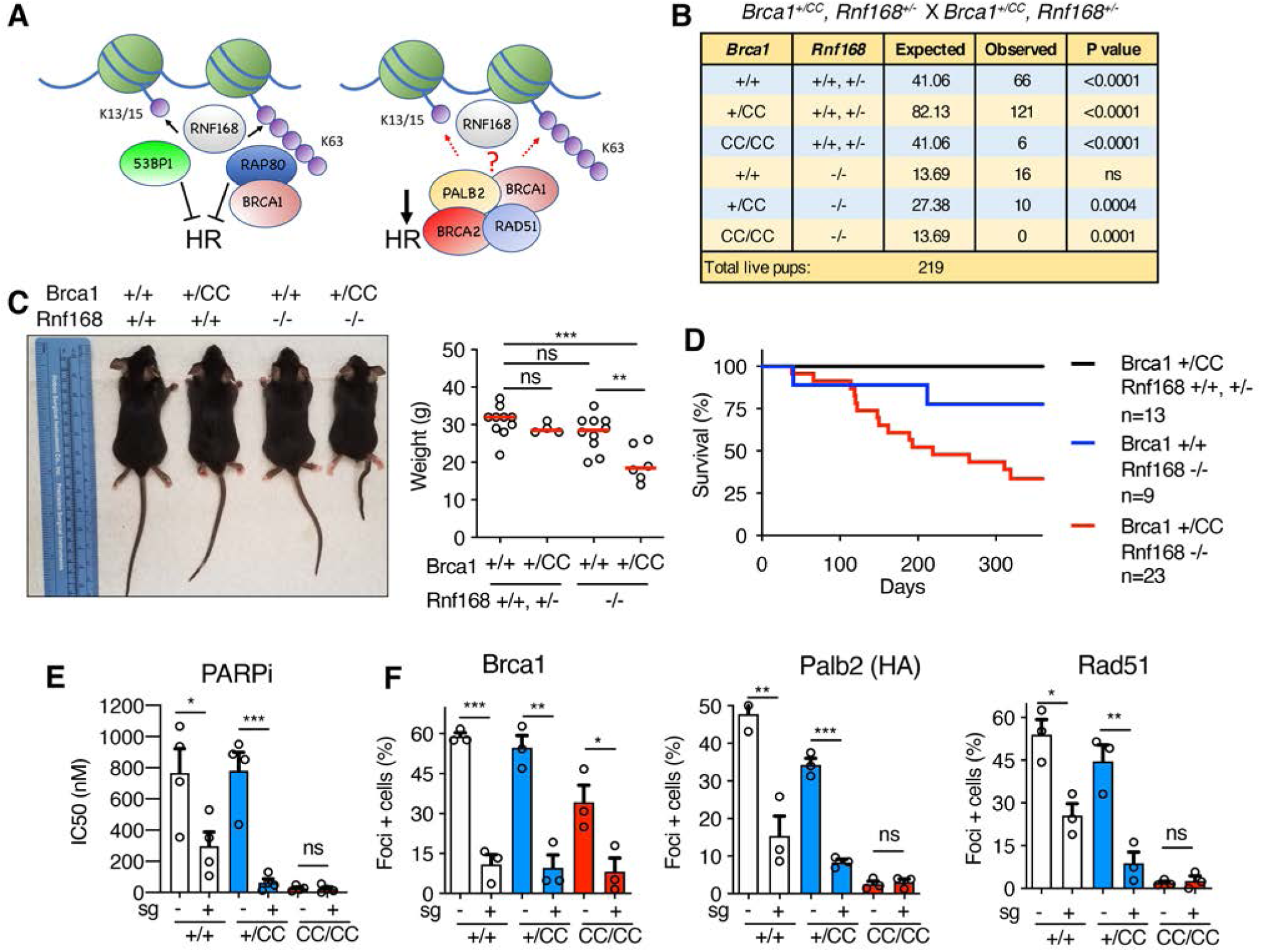
*Rnf168* and *Brca1* mutations block development and Brca1-P localization. **(A)** Cartoon showing RNF168-mediated and H2A-associated ubiquitin modifications, as well as the influence of 53BP1, BRCA1-A (ABRAXAS-RAP80) and -P (PALB2-BRCA2-RAD51) complexes on HR. **(B)** *Brca1^+/CC^, Rnf168^+/-^* mice were intercrossed and the numbers of expected and observed live births are shown. P value are obtained from chi-square goodness of fit tests for the binomial of each genotype. **(C)** *Left*; representative photograph. *Right*; weights of individual 4 to 6-month-old mice with the indicated genotypes, red bar - median. **(D)** Kaplan-Meier survival analysis of mice born with the indicated genotypes. **(E)** *Brca1^+/+^*, *Brca1^+/CC^*, and *Brca1^CC/CC^* MEFs with *GFP* (-) or *Rnf168* (+) targeting sgRNA were subsequently incubated with increasing concentrations of the PARPi rucaparib and colony formation assessed. Mean and S.E.M. rucaparib IC50 values are shown for n=4 biological replicates. **(F)** Cells from E were subject to 10 Gy γ-irradiation (IR) and foci formation measured by immunofluorescence. Endogenous Brca1 and Rad51 foci, and ectopic HA-PALB2 foci formation was assessed using cell lines expressing HA-PALB2. Foci positive cells were determined to be ≥10 Brca1 and Palb2 foci/nuclei, or ≥5 Rad51 foci/nuclei. The mean and S.E.M. percentage foci positive cells are shown for n=3 biological replicates. See **Fig. S2A** for representative images of cells. *** p < 0.001, ** p < 0.01, * p < 0.05, ^ns^ not significant (unpaired, two-tailed t-tests).

First, phenotypes associated with mice harboring *Brca1^CC^* and *Rnf168^-^* alleles were examined. The *Brca1^CC^* mutation specifically disrupts the Brca1-Palb2 interface^15^, and homozygous mice are born at reduced rates, show defective Rad51 loading and demonstrate developmental defects analogous to Fanconi anemia (FA). *Rnf168^-/-^* mice and cells displayed expected phenotypes (**Fig. S1A-C**), including defective 53bp1 foci formation and increased Rpa32 foci positive cells (**Fig. S1D**). Intercrossing *Brca1^+/CC^*, *Rnf168^+/-^* mice revealed that the *Brca1^CC/CC^; Rnf168^-/-^* genotype confers early embryonic lethality (**Fig. 1B** **and S1E**). Moreover, *Brca1^+/+^; Rnf168^-/-^* and *Brca1^+/CC^; Rnf168^+/+^* were found at the expected rates, but *Brca1^+/CC^, Rnf168^-/-^* mice were born at significantly reduced frequencies (**Fig. 1B**). Remarkably, *Brca1^+/CC^; Rnf168^-/-^* mice had traits that were similar to those observed in *Brca1^CC/CC^; Rnf168^+/+^* mice^15^, such as short stature and reduced weights (**Fig. 1C**), kinked tails, and areas of hypopigmented fur (**Fig. S1F**). *Brca1^+/CC^; Rnf168^-/-^* mice had reduced life spans (**Fig. 1D**), due to the onset of metastatic lymphoma (**Fig. S1G,H**). Additionally, assessment of PARPi sensitivity, which is indicative of HR-deficiency, showed that *Rnf168* knockout (KO) resulted in a 2.6- and 12.4-fold decrease in rucaparib LC50 values in *Brca1^+/+^* and *Brca1^+/CC^* MEFs, respectively. However, *Rnf168* KO did not affect *Brca1^CC/CC^* MEFs, which are highly sensitive from the outset (**Fig. 1E** and **Fig. S2A**). These results are in keeping with previous work showing that Rnf168 supports HR and development in mice with alleles that reduce the Brca1-Palb2 interaction^16^.

To determine whether mouse phenotypes were related to Brca1-P complex recruitment, we investigated the effects of *Rnf168* KO and the *Brca1* CC mutation on Brca1, Palb2, and Rad51 foci formation. As expected, *Rnf168* KO resulted in loss of 53bp1 IRIF positive cells (**Fig. S2A**). Of note, *Brca1^CC/CC^* cells produce a Brca1-CC protein that is stable and forms foci^15^. Strikingly, Brca1-WT and Brca1-CC IRIF were dramatically reduced in *Rnf168* in KO cells (**Fig. 1F** and **S2A,B**). The number of Palb2 and Rad51 IRIF positive *Brca1^+/+^* and *Brca1^+/CC^* cells also decreased in the absence of Rnf168. However, *Brca1^CC/CC^* cells, which have defective Brca1-Palb2 interaction, had few detectable Palb2 and Rad51 foci; therefore, *Rnf168* KO had little measurable impact (**Fig. 1F** and **S2A,B**). Overall, these results suggest a model whereby Rnf168 promotes Brca1 localization, and consequently, the Brca1-dependent recruitment of Palb2-Rad51 to DSBs.

RNF168-generated and K63-linked ubiquitin polymers are known to recruit RAP80^11–13^, and thereby the BRCA1-A complex to DSB sites. To determine if RAP80 is the mediator of BRCA1-P complex recruitment, we directly compared *Rap80* and *Rnf168* KO MEFs for Brca1 and Rad51 IRIF (**Fig. S2C**). In contrast to *Rnf168^-/-^* cells, there was a mild reduction in Brca1, and no effect on Rad51 foci positive *Rap80^-/-^* cells (**Fig. S2C**), suggesting that RNF168 recruits the BRCA1-P complex through another pathway.

BRCA1 DSB localization has previously been associated with both the RING and BRCT domains^12–14,17–19^. To explore mechanisms by which RNF168 promotes the recruitment of BRCA1 to DSBs, we assessed ectopic BRCA1 full-length (FL), BRCA1-ΔRING, and BRCA1-ΔBRCT foci formation (**Fig. 2A**). Cells expressing BRCA1-ΔBRCT had a moderate decrease, but BRCA1-ΔRING expressing cells had significantly fewer foci positive cells and foci per nucleus, compared with BRCA1-FL expressing cells (**Fig. 2B**). Interestingly, BRCA1-ΔRING also demonstrated a reduction in the overall size of foci (**Fig. 2B**). To examine whether protein expression impacted foci formation, we compared the BRCA1-ΔRING construct used above (aa 128-1863) to a more truncated construct that we previously found had improved expression and is produced from the endogenous *BRCA1^185delAG^* locus (aa 297-1863). Here, foci were similar between deletions, despite BRCA1-297-1863 showing higher expression (**Fig. S2D**).

**Figure 2.**
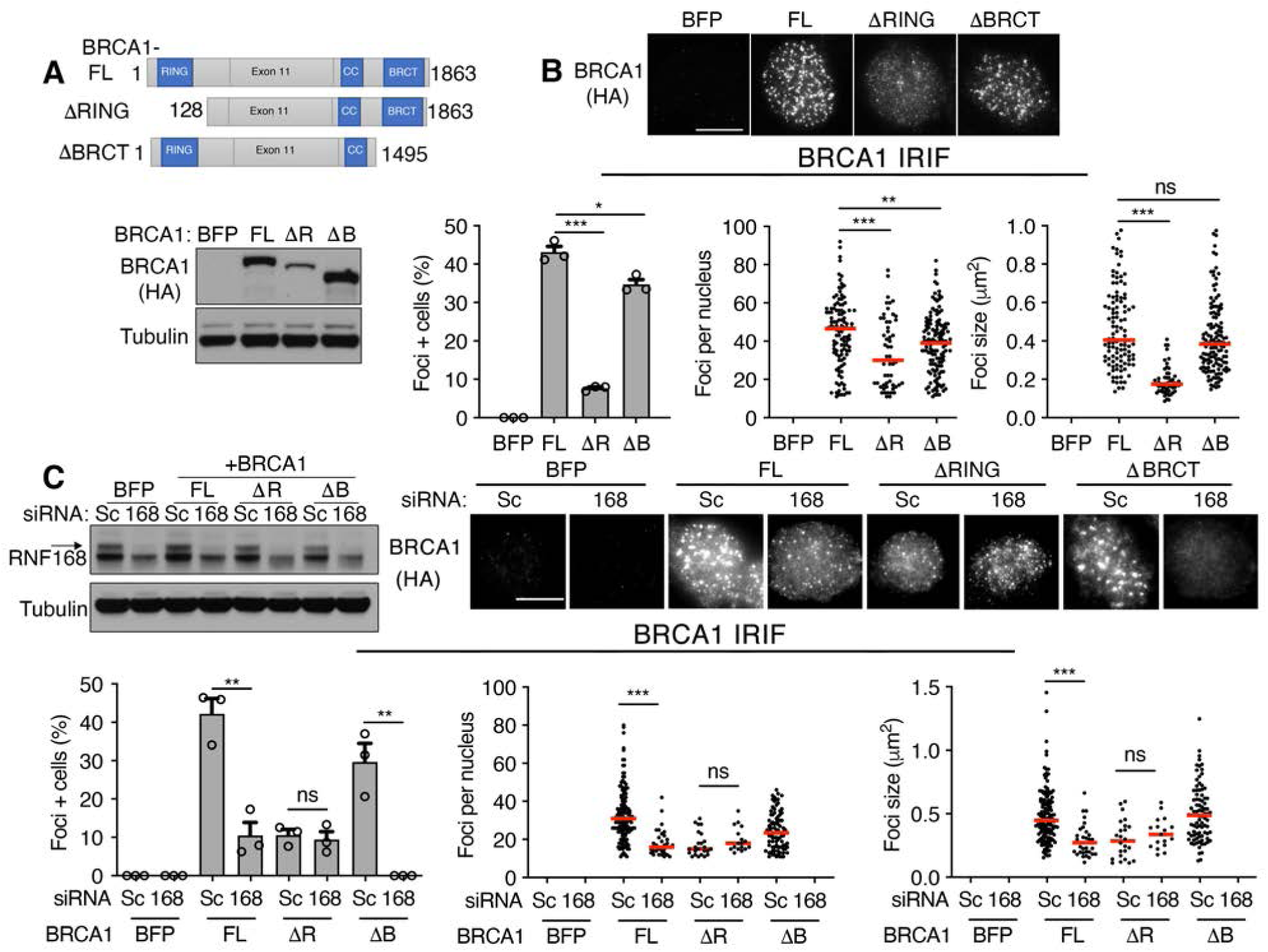
RNF168 regulates BRCA1 RING-mediated foci. **(A)** *Above*; cartoon showing BRCA1 protein domains and truncations with aa numbers indicated. *Below*; BRCA1 null MDA-MB-436 cells expressing BFP control, HA-BRCA1-full-length (FL), HA-BRCA1-ΔRING (ΔR), and HA-BRCA1-ΔBRCT (ΔB) were assessed for HA protein levels by Western blotting. **(B)** Cells from A were assessed for HA foci 8 hours post IR by immunofluorescence. *Above*; representative images (scale bar, 10 μm). *Left*; mean and S.E.M. percentage foci positive cells; *middle*, foci positive cells were assessed for the number of foci present in a single nucleus, red line: median value; *right*, foci positive cells were assessed for the mean size of foci present per nuclei, n=3 biological replicates. Foci positive cells were determined to be those with ≥10 foci/nuclei. *** p < 0.001, ** p < 0.01, * p < 0.05, ^ns^ not significant compared to FL (percentage positive cells: unpaired, two-tailed t-tests size; number and size per nucleus: nonparametric Mann-Whitney tests). **(C)** Cells from A were subject to scrambled (Sc) and RNF168-targeting siRNA followed by IR and HA foci formation measured by immunofluorescence. *Left*; Western blot showing RNF168 depletion. *Above*; representative cells (scale bar, 10 μm). *Below*; foci analyses as described in B, n=3 biological replicates.

We next measured the effects of RNF168 expression levels on BRCA1 foci formation. As expected, RNF168 siRNA reduced the number of foci positive cells as well as size of BRCA1-FL foci. Additionally, RNF168 siRNA resulted in a complete loss of detectable BRCA1-ΔBRCT foci; by contrast, BRCA1-ΔRING foci were unaffected (**Fig. 2C**). Thus, RNF168-induced BRCA1 focal accumulation requires the RING domain.

We aimed to decipher the mechanism by which the BRCA1 RING domain promotes foci formation. BRCA1 and BARD1 both contain N-terminal RING domains that directly interact and form an E3 ubiquitin ligase^20^. Because BRCA1-ΔRING does not interact with BARD1 nor engender ubiquitin ligase activity, we separately disrupted the E3 ligase activity versus the BRCA1-BARD1 interaction. The BRCA1-I26A mutation is commonly used to block ubiquitin ligase activity, but may retain residual activity. However, a triple I26A+L63A+K65A mutation (BRCA1-3A) fully abrogates E3 ligase activity, while retaining the ability to interact with BARD1^21^. The BRCA1-C61G protein retains the RING domain but prevents BRCA1-BARD1 heterodimerization and results in decreased protein stability^22^. However, we obtained cells with suitable ectopic protein expression (**Fig. 3A** and **S3A**). We found that BRCA1 foci formation was identical between BRCA1-FL and BRCA1-3A expressing cells. By contrast, BRCA1-C61G formed fewer foci that were smaller and similar in size to the BRCA1-ΔRING protein (**Fig. 3B**). Similar to BRCA1-ΔRING, RNF168 siRNA also had no effect on BRCA1-C61G foci (**Fig. S3B**). To further ensure that E3 ligase activity was absent, we generated a construct that physically linked BRCA1-ΔRING and BARD1-ΔRING peptides (BRCA1-ΔRING-L-BARD1-ΔRING), thereby creating a synthetic heterodimer where both proteins were devoid of RING domains (**Fig. 3A** and **S3A**). Here, BRCA1-ΔRING-L-BARD1-ΔRING IRIF was identical to BRCA1-FL (**Fig. 3B**), suggesting that the interaction with BARD1 is required for efficient BRCA1 focal accumulation.

**Figure 3.**
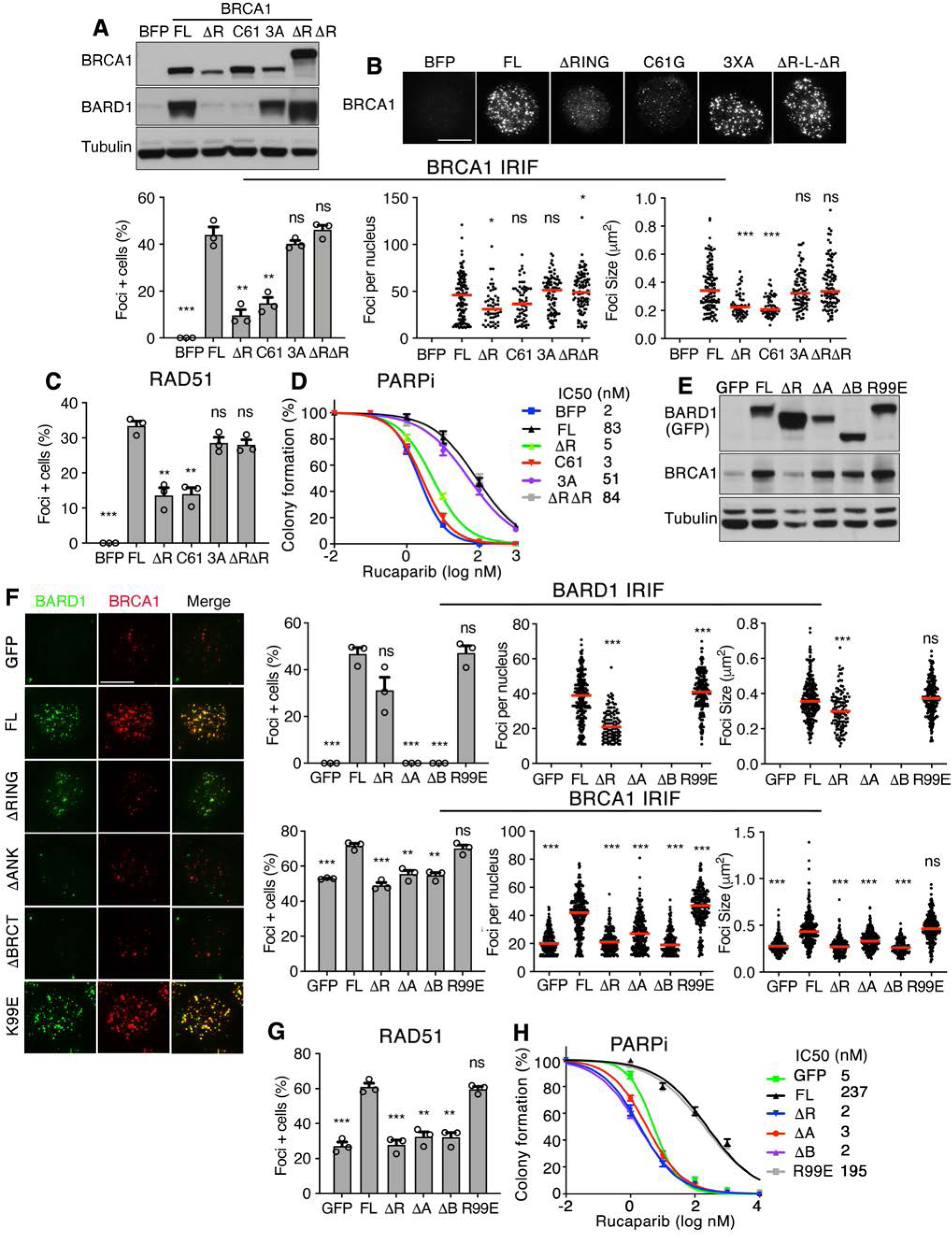
BARD1 ANK and BRCT domains promote BRCA1 foci formation. **(A)** MDA-MB-436 cells expressing BFP, BRCA1-full-length (FL), BRCA1-ΔRING (ΔR), BRCA1-C61G (C61), BRCA1-I26A+L63A+K65A (3A), BRCA1-ΔRING-L-BARD1-ΔRING (ΔRΔR) were assessed for BRCA1 and BARD1 expression by Western blotting. **(B)** Cells from A were assessed for BRCA1 foci 8 hours post IR by immunofluorescence. *Above*; representative images (scale bar, 10 μm). *Left*; mean and S.E.M. percentage foci positive cells; *middle*, foci positive cells were assessed for the number of foci present in a single nucleus, red line: median value; *right*, foci positive cells were assessed for the mean size of foci present per nuclei, n=3 biological replicates. Foci positive cells were determined to be those with ≥10 foci/nuclei. *** p < 0.001, ** p < 0.01, * p < 0.05, ^ns^ not significant (percentage positive cells: unpaired, two-tailed t-tests size; number and size per nucleus: nonparametric Mann-Whitney tests). **(C)** Cells from A were assessed for RAD51 foci 8 hours post IR by immunofluorescence. See **Fig. S3C** for representative images. Mean and S.E M. percentage (≥5) foci positive cells, n=3 biological replicates. *** p < 0.001, ** p < 0.01, ^ns^ not significant compared to FL (unpaired, two-tailed t-tests). **(D)** Cells from A were incubated with increasing concentrations of rucaparib for 2 weeks and colony formation assessed. Mean and S.E.M. colony formation as well as mean IC50 values, n=3 biological replicates. **(E)** MCF7 *BARD1^-/-^* cells expressing GFP control, GFP-BARD1-full-length (FL), GFP-BARD1-ΔRING (ΔR), GFP-BARD1-ΔANK (ΔA), GFP-BARD1-ΔBRCT (ΔB), and GFP-BARD1-R99E (R99E) were assessed for GFP and BRCA1 expression by Western blotting. **(F)** Cells from E were assessed for BARD1 and BRCA1 foci 8 hours post IR by immunofluorescence. *Left*; representative foci (scale bar, 10 μm). *Right*; foci analyses as described in B. **(G)** Cells from E were assessed for RAD51 foci as described in C. See **Fig. S3H** for representative images. **(H)** Cells from E were incubated with increasing concentrations of rucaparib for 2 weeks and colony formation assessed. Mean and S.E.M. colony formation as well as mean IC50 values, n=3 biological replicates.

Interestingly, while BRCA1-FL, BRCA1-3A, and BRCA1-ΔRING-L-BARD1-ΔRING expressing cells showed increased RAD51 IRIF, BRCA1-C61G and BRCA1-ΔRING expressing cells had significantly lower levels of RAD51 IRIF (**Fig. 3C** and **S3C**). PARPi sensitivity is reflective of cellular HR proficiency, and using standard colony assays, BRCA1-FL, BRCA1-3A, and BRCA1-ΔRING-L-BARD1-ΔRING expressing cells were resistant, but BRCA1-C61G and BRCA1-ΔRING expressing cells PARPi sensitive (**Fig. 3D**). However, we confirmed previous results using PARPi resistance colony assays^23^, with BRCA1-C61G and BRCA1-ΔRING showing hypomorphic activity and providing residual PARPi resistance relative to BFP-expressing control cells (**Fig. S3D**).

The above results indicate that BARD1 recruits BRCA1, and consequently RAD51 to DNA damage sites. In line with the known requirement of BARD1 in providing BRCA1 protein stability^24^, CRISPR/Cas9-mediated KO of *BARD1* reduced endogenous BRCA1 protein expression levels (**Fig. S3E**), as well as the number of BRCA1 and RAD51 foci in MCF7 cells (**Fig. S3F**). To identify regions of BARD1 critical for BRCA1 recruitment, we expressed ectopic BARD1-FL, BARD1-ΔRING, BARD1-ΔANK and BARD1-ΔBRCT truncated proteins, as well as an E3 ligase disrupting BARD1-R99E mutation^25^ (**Fig. 3E** and **S3G**). Of note, all ectopic BARD1 mutations except for BARD1-ΔRING restored endogenous BRCA1 protein expression (**Fig. 3E**). Using pre-extraction methods, BARD1-FL and BARD1-R99E formed similar, and BARD1-ΔRING less efficient, foci. However, BARD1-ΔANK and BARD1-ΔBRCT foci formation was barely detectable (**Fig. 3F**). The number of BRCA1 foci per nucleus and size of foci were markedly reduced in BARD1 KO cells, but were restored in BARD1-FL and BARD1-R99E expressing cells. In contrast, there remained fewer and smaller BRCA1 foci in BARD1-ΔRING, BARD1-ΔANK and BARD1-ΔBRCT expressing cells (**Fig. 3F**). RPA32 and RAD51 foci formation followed similar patterns as BRCA1 in ectopic add back cells (**Fig. 3G** and **S3H**). PARPi resistance was also restored in BARD1-FL and BARD1-R99E, but not BARD1-ΔRING, BARD1-ΔANK and BARD1-ΔBRCT expressing cells (**Fig. 3H**). These results establish the BRCA1-BARD1 interaction, as well as the BARD1 ANK and BRCT domains, as mediators of BRCA1 focal accumulation.

A recent study revealed that BARD1 localizes to DNA damage sites through a dual mechanism involving both the ANK and BRCT domains^26^. The BARD1 ANK domain is a reader of histone H4 unmethylated at K20 (H4K20me0)^17^, and the BRCT repeat was shown to contain a BRCT domain ubiquitin-dependent recruitment motif (BUDR), which directly binds mUb-H2A^26^. Indeed, MCF7 *BARD1^-/-^* cells engineered to express a *BARD1^BUDR^* mutant construct (**Fig. S4A**), had identical phenotypes to *BARD1^ΔBRCT^* expressing cells, with inefficient BARD1 DSB localization (**Fig. 4A** and **Fig. S4B**), and failed to rescue BRCA1 and RAD51 foci formation (**Fig. 4A, B** and **S4B**), nor PARPi sensitivity (**Fig. 4C**).

**Figure 4.**
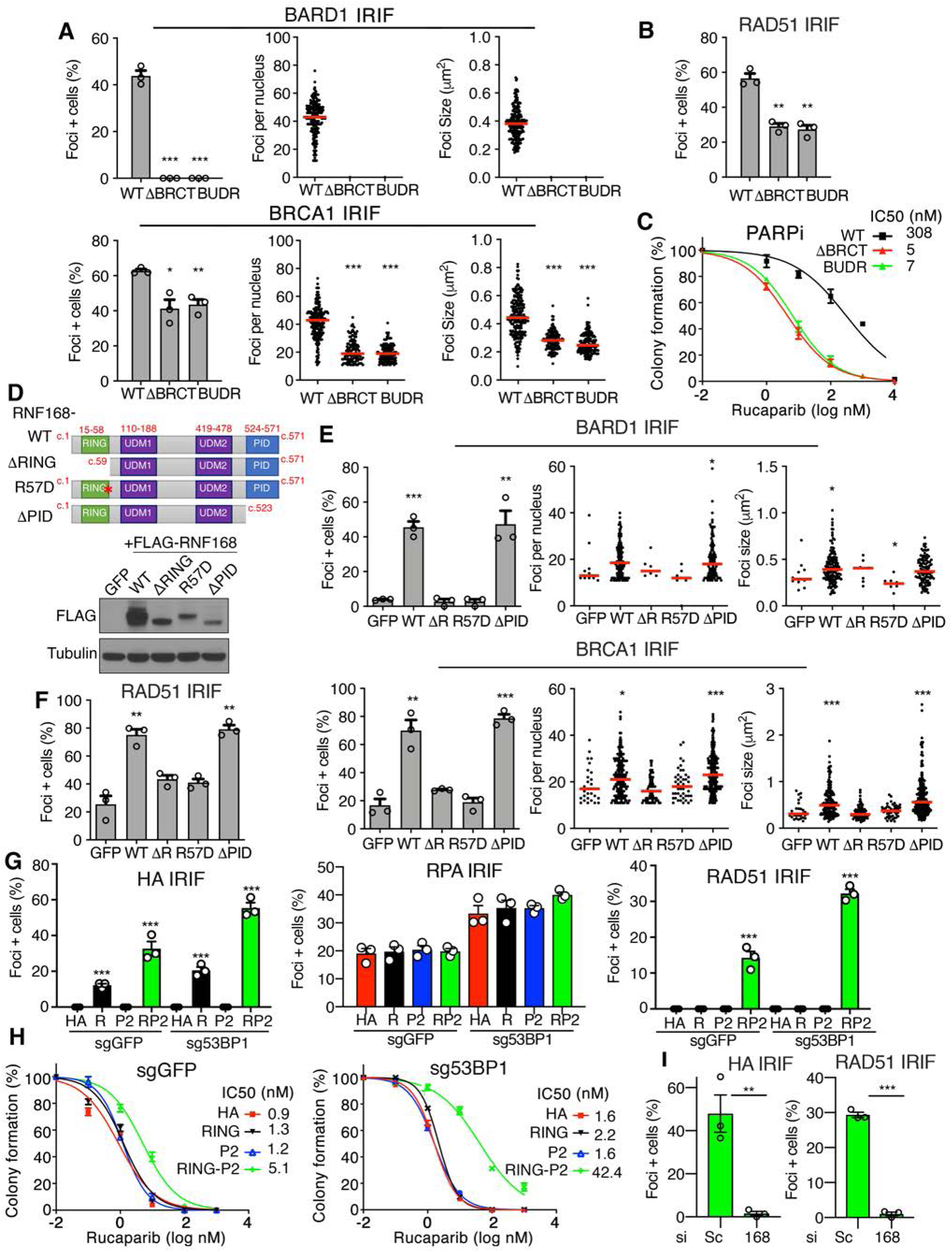
RNF168-mediated H2A mono-ubiquitination recruits BARD1 via the BUDR. **(A)** MCF7 *BARD1^-/-^* cells expressing GFP-BARD1-WT, GFP-BARD1-ΔBRCT, or GFP-BARD1-BUDR (R705A+D712A+Q715R) were assessed for BARD1 and BRCA1 foci size, number per nucleus, as well percentage foci positive cells post IR by immunofluorescence. See **Fig. S4B** for representative images. *Left*; mean and S.E.M. percentage foci positive cells; *middle*, foci positive cells were assessed for the number of foci present in a single nucleus, red line: median value; *right*, foci positive cells were assessed for the mean size of foci present per nuclei, n=3 biological replicates. Foci positive cells were determined to be those with ≥10 foci/nuclei. *** p < 0.001, ** p < 0.01, * p < 0.05, ^ns^ not significant compared to WT (percentage positive cells: unpaired, two-tailed t-tests size; number and size per nucleus: nonparametric Mann-Whitney tests). **(B)** Cells from A were assessed for RAD51 foci 8 hours post IR by immunofluorescence. Mean and S.E M. percentage (≥5) foci positive cells, n=3 biological replicates. ** p < 0.01 compared to WT (unpaired, two-tailed t-tests). See **Fig. S4B** for representative images. **(C)** Cells from A were incubated with increasing concentrations of rucaparib for 2 weeks and colony formation assessed. Mean and S.E.M. colony formation as well as mean IC50 values, n=3 biological replicates. **(D)** Cartoon of RNF168 protein domains and mutations analyzed. *Below*; expression of ectopic FLAG-RNF168 proteins by Western blotting in MDA-MB-231 *RNF168^-/-^* cells. **(E)** Cells from D were assessed for BARD1 and BRCA1 foci as described in A with statistical comparisons to cells expressing GFP. See **Fig. S4G** for representative images. **(F)** Cells from D were assessed for RAD51 IRIF as described in B with statistical comparisons to cells expressing GFP. See **Fig. S4G** for representative images. **(G)** MDA-MB-436 cells with sgRNA targeting GFP or 53BP1 were infected with virus expressing HA-RING, HA-PALB2, or HA-RING-PALB2 and assessed for HA (≥10), RPA32 (≥10), and RAD51 (≥5) IRIF positive cells. Mean and S.E M. percentage foci positive cells, n=3 biological replicates. *** p < 0.001 compared to HA alone (unpaired, two-tailed t-tests). See **Fig. S5A,B** for western blots and representative images. **(H)** Cells from G were incubated with increasing concentrations of rucaparib for 2 weeks and colony formation assessed. Mean and S.E.M. colony formation as well as mean IC50 values are shown from n=3 biological replicates. **(I)** MDA-MB-436 cells with sgRNA targeting 53BP1 expressing HA-RING-PALB2 were subject to scrambled (Sc) or RNF168 siRNA and HA and RAD51 IRIF measured. Mean and S.E M. percentage foci positive cells, n=3 biological replicates. *** p < 0.001 and ** p < 0.01 compared to Sc (unpaired, two-tailed t-tests). See **Fig. S5C** for Western blots and representative images.

Given the established role of RNF168 E3 ligase activity in generating mUb-H2A^10^, we reasoned that RNF168 promotes BRCA1-P recruitment to DSB sites via BARD1 BUDR binding to mUb-H2A. To further investigate this possibility, we assessed a series of ectopic add back *RNF168* mutations using an MDA-MB-231 *RNF168^-/-^* clone (**Fig. S4C, D**). Similar to results in MEFs, BRCA1 and RAD51 IRIF were significantly reduced in MDA-MB-231 *RNF168^-/-^* cells (**Fig. S4E**). The *RNF168^ΔRING^* mutation results in loss of the RING domain and E3 ligase activity, whereas *RNF168^R57D^* retains E3 ligase activity but specifically blocks the ability of RNF168 to mono-ubiquitinate H2A^10^. Additionally, *RNF168^ΔPID^* lacks the C-terminally located PALB2-interacting domain (PID), which is required for RNF168 to directly interact with PALB2 and facilitate the latter’s foci formation in the absence of BRCA1^27^ (**Fig. 4D**). The mRNA expression of ectopic constructs was within a similar range, but RING domain mutant protein levels were dramatically reduced compared to wild-type (**Fig. S4F**), indicating that RING mutations are detrimental to protein stability. To bypass this issue, we used a *CMV* promoter for RING mutant constructs, which produced cell lines with comparable protein expression to *UBC* promoter-driven *RNF168^WT^* and *RNF168^ΔPID^* expressing cells (**Fig. 4D**). IRIF experiments revealed that RNF168^WT^ and RNF168^ΔPID^ formed robust foci and fully rescued 53BP1 IRIF. In contrast, RNF168^ΔRING^ and RNF168^R57D^ formed smaller foci and failed to rescue 53BP1 IRIF (**Fig. S4G**). Moreover, while RNF168^WT^ and RNF168^ΔPID^ rescued, RNF168^ΔRING^ and RNF168^R57D^ had no effect on BARD1 and BRCA1 (**Fig. 4E** and **S4G**), or RAD51 foci (**Fig. 4F** and **S4G**). These data indicate that the E3 ligase activity, and specifically the ability of RNF168 to generate mUb-H2A, is required for efficient BARD1-BRCA1-RAD51 DSB localization.

We postulated that in the absence of BRCA1, a direct association of only the BRCA1 RING domain with PALB2 may direct PALB2-RAD51 to DSBs and rescue HR. Thus, we generated a construct containing the BRCA1 RING domain linked to PALB2 cDNA (HA-RING-L-PALB2) and expressed in BRCA1 null MDA-MB-436 cells. RING-containing constructs increased BARD1 protein similar to endogenous levels observed in *BRCA1* wild-type MCF7 cells (**Fig. S5A**). Because MDA-MB-436 cells are BRCA1 null, and end resection is necessary for efficient PALB2-RAD51 loading^28^, we also subject cells to GFP- or 53BP1-targeting sgRNA, which increased RPA32 IRIF (**Fig. 4G** and **Fig. S5A**). HA-RING-L-PALB2 formed detectable foci in sgGFP cells, but HA-RING-L-PALB2 and RAD51 foci were dramatically increased (**Fig. 4G** and **Fig. S5B**), and PARPi resistance was observed in sg53BP1 cells (**Fig. 4H**). Moreover, we confirmed that HA-RING-L-PALB2 and RAD51 foci were entirely reliant on RNF168 activity (**Fig. 4I** and **Fig. S5C**). These data establish the BRCA1 RING domain as an RNF168-dependent DSB localization module.

Despite numerous pathogenic *BRCA1* mutations being located within the BRCT repeats, loss of the BRCT domain had relatively mild effects on BRCA1 protein localization (**Fig. 2B**). To gain additional insight into how BRCA1 BRCT domain mutations impact BRCA1 localization and HR, we compared several truncating mutations located at varying distances from the BRCT repeat region, which spans ∼aa’s 1642-1856 (**Fig. 5A**). We confirmed previous observations^29^, with stop codons located close to (aa 1558*), or within (aa 1814*), the BRCT domain, despite similar mRNA levels resulted in reduced protein expression (**Fig. 5A**), and consequently few foci forming cells (**Fig. 5B** and **S5D**). However, the BRCA1-ΔBRCT protein (aa 1495*), which stops ∼147 aa prior to the first BRCT repeat, was stable and formed relatively robust foci (**Fig. 5A,B**). Despite this, BRCA1-ΔBRCT expressing cells failed to promote CtIP, RPA32, and RAD51 IRIF, relative to full-length BRCA1 expressing cells (**Fig. 5B** and **S5D**). Thus, even when BRCA1 is stable and can localize to DSBs, in the absence of the BRCT domain, CtIP recruitment, and consequently end resection and HR, is inefficient.

**Figure 5.**
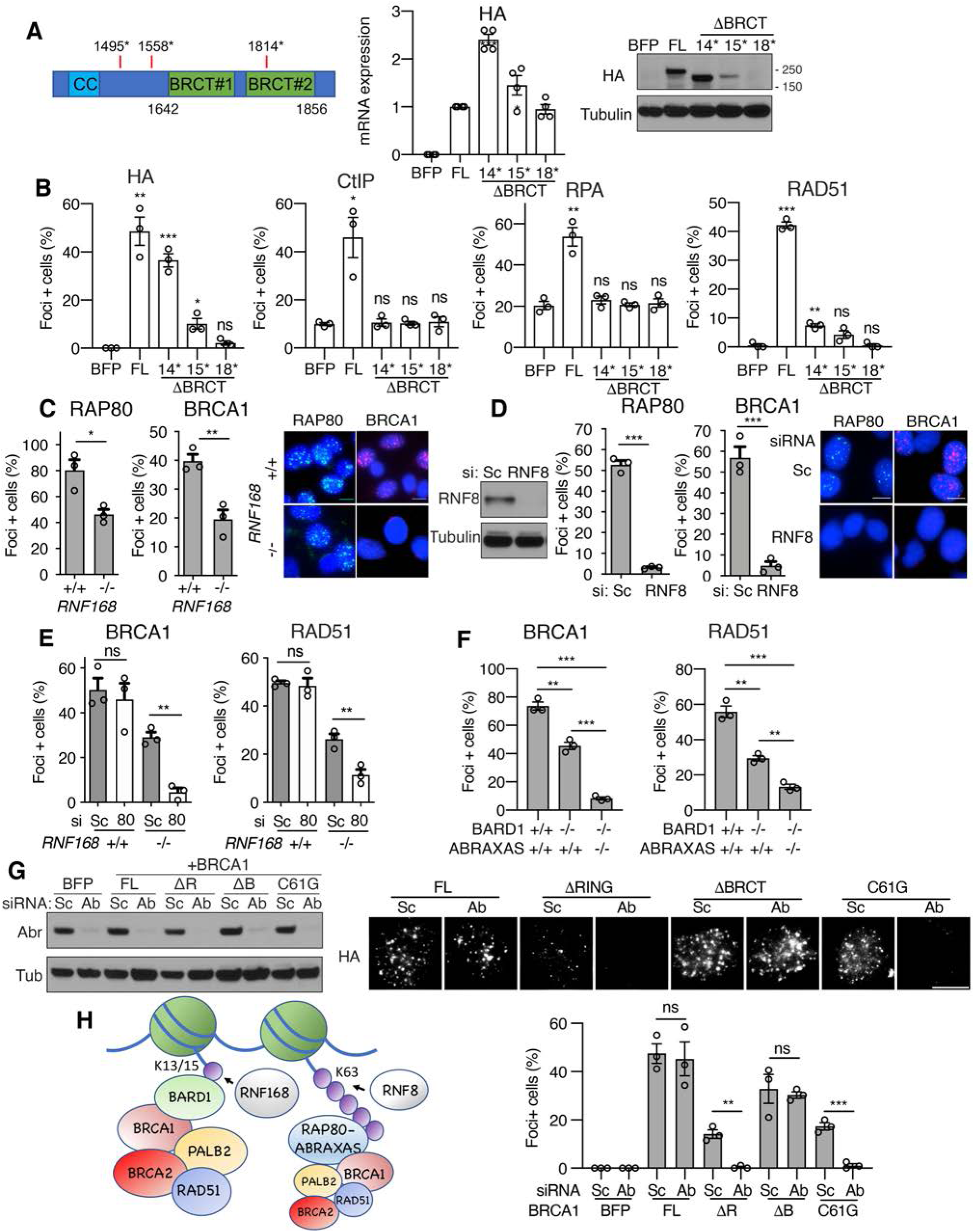
BRCA1 BRCT-ABRAXAS-RAP80 axis is a backup pathway. **(A)** *Left*; cartoon showing BRCA1 BRCT region with aa numbers indicated and positions of stop codon mutations tested. *Middle*; BRCA1 null MDA-MB-436 cells expressing BFP control, HA-BRCA1-full-length (FL), HA-BRCA1-1495stop (*), HA-BRCA1-1558*, and HA-BRCA1-1814* were assessed for HA mRNA and divided by a house-keeping gene expression by qRT-PCR, followed by normalization to HA-BRCA1-FL. *Right*; cells were assessed for HA protein levels by Western blotting. **(B)** Cells from A were assessed for HA (≥10), CtIP (≥10), RPA32 (≥10), and RAD51 (≥5) IRIF, with the indicated number of foci per nuclei counted as positive. See **Fig. S5D** for representative images. **(C)** MDA-MB-231 *RNF168*^+/+^ and *RNF168*^-/-^ cells were examined for RAP80 (≥10) and BRCA1 (≥10) IRIF-positive cells, inset representative images. **(D)** MDA-MB-231 *RNF168*^+/+^ cells were examined for RAP80 (≥10) and BRCA1 (≥10) IRIF-positive cells after Sc or RNF8 siRNA, inset representative images. **(E)** Cells from C were subject to scrambled (Sc) or RAP80 (80) siRNA treatment and assessed for BRCA1 (≥10) and RAD51 (≥5) IRIF-positive nuclei. See **Fig. S5E** for representative images and RAP80 western blot. **(F)** MCF7 *WT*, *BARD1*^-/-^, and *BARD1*^-/-^, *ABRAXAS*^-/-^ DKO cells were examined for BRCA1 and RAD51 IRIF positive cells. See **Fig. S5F** for representative images. **(G)** MDA-MB-436 cells expressing BFP, BRCA1-full-length (FL), BRCA1-ΔRING (ΔR), BRCA1-ΔBRCT (ΔB), or BRCA1-C61G (C61), were subject to Sc or ABRAXAS siRNA and assessed for BRCA1 foci formation. Mean and S.E.M. percentage foci positive cells from 3 biological replicates are shown throughout. *** p < 0.001, ** p < 0.01, * p < 0.05, ^ns^ not significant (unpaired, t-tailed t-tests). **(H)** Model for mechanisms of RNF168-mediated BRCA1-P complex recruitment to DSBs. BARD1 BUDR binding to mUb-H2A represents the dominant recruitment pathway, with RAP80-ABRAXAS contributing to a backup pathway.

Although loss of the BRCA1 BRCT domain or RAP80 induced mild defects in BRCA1 foci (**Fig. 2A** and **S2C**), we hypothesized that the BRCA1-A complex was likely important for residual BRCA1-P localization in the absence of the RNF168-BARD1-RING recruitment axis. Interestingly, RAP80 foci were reduced but detectable in *RNF168^-/-^* MDA-MB-231 cells, potentially accounting for the residual BRCA1 foci observed (**Fig. 5C**). In contrast, depletion of RNF8, which generates K63-linked ubiquitin chains that are required for both RNF168 and RAP80 recruitment^12–14,30^, resulted in a severe reduction in both RAP80 foci and BRCA1 foci (**Fig. 5D**). Therefore, the RNF8-BRCA1-A pathway appears to be partially active in the absence of RNF168. In support of this, RAP80 depletion dramatically reduced BRCA1 and RAD51 IRIF in *RNF168^-/-^* cells (**Fig. 5E** and **S5E**). Moreover, double knockout (DKO) *BARD1^-/-^*, *ABRAXAS^-/-^* MCF7 cells also had severely impaired BRCA1 and RAD51 foci relative to single KO *BARD1^-/-^* cells (**Fig. 5F** and **S5F**), confirming the supporting role of the BRCA1-A complex. Last, ABRAXAS RNAi had no effect on cells expressing BRCA1-full-length or BRCA1-ΔBRCT proteins, but completely abrogated BRCA1-ΔRING and BRCA-C61G IRIF (**Fig. 5G**). Thus, the BRCA1-A complex is likely critical for residual HR activity in cancers with *BRCA1* RING domain located mutations. All in, the recruitment of the BRCA1-P complex to DSBs occurs primarily through an RNF168-mUb-H2A-BARD1-BRCA1 RING axis, with RNF8-RAP80-ABRAXAS-BRCA1 BRCT signaling providing a backup route (**Fig. 5H**).

In the current study, we report an RNF168-governed pathway that is responsible for the recruitment of the BRCA1-P complex to DSBs. A BUDR was recently discovered within the BARD1 BRCT domain, which directly binds mUb-H2A^26^. In support of this, mutating residues within the BUDR, or specifically blocking the ability of RNF168 to generate mUb-H2A, reduced BRCA1-P complex recruitment to DSBs.

We found human and mouse cell lines to be highly dependent on RNF168 activity for efficient BRCA1, BARD1, and RAD51 foci formation. Loss of RNF168 activity has previously been shown to reduce BRCA1 foci formation^9,11,27^. However, others reported mild or little impact^16,31^, and we are unable to pinpoint reasons for these discrepancies, but speculate assessment of foci versus laser-stripes, or quantification methods, could contribute. RNF168 is capable of promoting BRCA1-A complex recruitment through the production of K63-linked ubiquitin polymers that form a binding module for RAP80^11,13^. Moreover, RNF8 generated K63-linked ubiquitin chains^6,32–34^, as well as SUMO-binding^35–37^, can contribute to RAP80 foci. In agreement, *RNF168^-/-^* cells had reduced, but detectable RAP80 foci. Interestingly, BRCA1-A complex disruption was highly detrimental for BRCA1-P activity in *RNF168^-/-^* and *BARD1^-/-^* cells, suggesting the RAP80-ABRAXAS axis confers a back-up recruitment pathway (**Fig. 5H**). Importantly, individual disruption of the RING or BRCT recruitment pathways were insufficient to entirely abolish BRCA1 foci, shedding light on how *BRCA1* mutant cancers utilize hypomorphic proteins, which often lack these domains, to contribute to residual HR activity and PARPi resistance^23,29, 38–40^.

Although the BRCA1-A complex provided inefficient BRCA1-P localization, it is likely sufficient to maintain residual HR and the relatively normal development of *Rnf168*^-/-^ mice. *Brca1^+/CC^* reduces the number of intact Brca1-Palb2 complexes, and *Brca1^+/CC^*, *Rnf168*^-/-^ KO mice had a greater reduction in Palb2 and Rad51 foci compared to either genotype alone, as well as similar developmental defects to *Brca1^CC/CC^*, *Rnf168*^+/+^ mice^15^. Moreover, while we were unable to detect Palb2 and Rad51 foci in *Brca1^CC/CC^* cells, it is possible that residual Rad51 loading, below the foci detection threshold, account for the birth of *Brca1^CC/CC^* live mice. In contrast, *Brca1^CC/CC^*, *Rnf168*^-/-^ resulted in early embryonic lethality, potentially due to a complete abolishment of residual Rad51 loading (**Fig. S5G**). Another possibility is that Rnf168 compensates for loss of Brca1 activity and directly loads Palb2^27^. However, we were unable to detect Palb2 or Rad51 foci in *Rnf168^+/+^*, *Brca1^CC/CC^* cells, making the role of the Rnf168-Palb2 interface unclear in this setting.

Despite both *Rnf168* and *53bp1* KO activating resection, the effects on *Brca1* mutant mouse viability differ due to the additional requirement of Rnf168 in recruiting Brca1, and subsequently Palb2-Rad51 to DSBs. HR and developmental defects associated with mice harboring *Brca1* exon 2 mutations were previously shown to be rescued by *Rnf168* KO due to loss of 53bp1 localization and DNA end resection activation^16^. *Brca1* exon 2 mutations produce a Brca1 Ringless protein isoform, resulting in loss of Bard1 interaction^41^. Here, Rnf168 is dispensable for Rad51 foci and HR, possibly due to loss of the Brca1 Ring domain and an inability to influence Bard1-mediated Brca1-P recruitment. Thus, the BRCA1 RING domain could determine phenotypes associated with modulation of RNF168 activity in cancer. *BRCA1 185delAG* and C61G mutations are commonly found in patients and produce proteins that fail to interact with BARD1^22,23,39,42^. Cancers with these mutations may reduce RNF168 activity as a means of promoting DNA end resection and inducing RAP80-mediated RINGless BRCA1-P loading, and consequently PARPi resistance. In contrast, hypomorphic proteins generated that retain the RING, may be highly dependent on RNF168 for residual BRCA1 protein activity^43^.

BARD1 and 53BP1 both bind the same mUb-H2A modification^26^, implicating protein occlusion as a DSB resection regulatory mechanism. However, the switch to resection is likely more complex, given BRCA1 exon 11 and BRCT regions are also required for efficient resection^15,16,44,45^. Indeed, the BRCA1 RING peptide was capable of stabilizing BARD1 and localizing to DSBs, but insufficient to activate resection (**Fig. 4G**). Overall, we establish a series of molecular interactions linking RNF168-ubiquitin signaling to BRCA1-PALB2-associated HR. Intriguingly, BRCA1 molecules may simultaneously localize to DSBs, activate resection, and promote PALB2-RAD51 loading.

## Methods

### Mouse models

The Fox Chase Cancer Center (FCCC) Institutional Animal Care and Use Committee (IACUC) approved experiments involving mice. Generation of mice harboring the *Brca1^CC^* allele was previously described^15^. *Rnf168*^-^ mice were generated with the help of the FCCC Transgenic Mouse facility using CRISPR/Cas9 technology. Guide RNA sequences targeting the *Rnf168* Ring domain were selected based on high predicted cut efficiency and minimal predicted off-target sites (https://portals.broadinstitute.org/gpp/public/analysis-tools/sgrna-design). Several sgRNAs were tested and ultimately the sgRNA target sequence TCTGTCGTCGCCGGGTCTCT was used. Cas9 enzyme (IDT, 1081058) and 20 ng/μl sgRNA were resuspended in injection buffer (1 mM Tris-HCl pH 7.5, 0.1 mM EDTA) and injected into single cell embryos in a B6C3F1/J background (offspring of C57BL/6J females and C3H/HeJ males, Jax Laboratory). Mice derived from injected embryos were subsequently crossed with C57BL/6J mice, resulting in a mixed genetic background. Developmental abnormalities in mice such as tail kinks and hypopigmentation were documented after visual inspection. Kaplan-Meier survival was assessed upon mice reaching IACUC guidelines for euthanasia and organs collected, fixed in neutral buffered formalin, paraffin embedded, sections hematoxylin and eosin stained and evaluated by the FCCC Histopathology facility. T-cell lymphoma diagnoses were confirmed by immunohistochemical staining of sections with anti-CD3 antibody (Agilent, M725429-2) and DAB solution development.

### Mouse genotyping, embryo analysis, MEF generation and culture

DNA from mice were extracted with 240 μl 50 mM NaOH followed by a 10-minute incubation at 95 degrees then neutralized with 60 μl 1 M Tris-HCl, pH 8. Genotyping for *Brca1^CC^* status was accomplished by PCR amplification using primers (GGTGCACTCTCCTCCAACATC, CTCTTGCACCTGCCTCTCTGA). PCR products were subject to digestion with EcoNI and bands visualized using agarose gel electrophoresis. The *Brca1^CC^* allele introduces an additional EcoNI cut site, as previously described^15^. *Rnf168^-^* allele was detected using two PCR reactions with a common primer (GGGTCCTCACCGCGTAAGAAG) and one reaction that amplifies the wild-type gene (CGAAGAGACCCGGCGACGA) and another that amplifies the mutant allele (GTATGGTACCGAGTCCATAAGGGACA) and bands visualized using agarose gel electrophoresis. Pregnant mice were euthanized at 12.5 and 13.5 days post-coitum for assessment of Mendelian developmental ratios and generation of mouse embryonic fibroblast (MEF) cell lines, respectively. Embryos were separated and collected for genotyping. To generate MEF lines, individual embryos were washed with sterile Dulbecco’s phosphate-buffered saline (DPBS). Embryos were aspirated 3 times through a 16 G needle and incubated for 15 min in trypsin/EDTA at 37 degrees. Cells were pelleted by centrifugation for 5 min at 1000g, resuspended in 10 ml MEF media (Dulbecco’s Modified Eagle’s Medium with 4.5 g/L glucose, 15% fetal bovine serum, 1x GlutaMAX, 1x MEM nonessential amino acids, 1 mM sodium pyruvate, 1x penicillin/streptomycin). Prior to passage three, MEFs were immortalized with pBabe-SV40Tag by electroporation using the Amaxa Mouse/Rat Hepatocyte Nucleofector Kit (Lonza, VPL-1004). Immortalized MEF lines were genotyped, tested for mycoplasma and maintained in MEF media with 10% FBS.

### Cell lines

MDA-MB-436, MDA-MB-231 and MCF7 cell lines were obtained from ATCC, tested for mycoplasma (Lonza, LT07-705) and identities confirmed by short tandem repeat (STR) profiling using IDEXX analysis. Cells were grown at 37 degrees with 5% CO_2_ in RPMI media with 10% FBS and 1x penicillin/streptomycin.

### CRISPR/Cas9 edited cell lines

MEF cell lines were transduced with lentivirus containing Cas9 and guide RNA in the lentiCRISPRv2 plasmid (Addgene, 52961) and selected using 2 μg/ml puromycin. Single guide RNA (sgRNA) target sequences were selected based on low predicted off-target score and predicted cutting efficiencies (https://portals.broadinstitute.org/gpp/public/analysis-tools/sgrna-design) then cloned using BsmBI sites. The *Rnf168*-targeted guide utilized the sequence TCTGTCGTCGCCGGGTCTCT to generate MEF lines. Single cell clones were isolated where indicated and were considered a knockout (KO) if all alleles exhibited frameshifting mutations by Sanger sequencing using the primers described for mouse genotyping. KO was further confirmed by the absence of 53bp1 IRIF. *Rap80* was targeted with ATAGTGATATCCGATAGCGA sgRNA target sequence and KO confirmed by Sanger sequencing using the following primers; GGTAAAAGGATGCCACGAAGGA and CAAATAATATGACCACACGCGCT. MDA-MB-231 CRISPR/Cas9 edited cell lines were generated by first isolating clones with doxycycline-inducible expression of Cas9 (Addgene, 50661), then infected with lentivirus harboring the sgRNA and selected with 4 μg/ml blasticidin. Surviving cells were subjected to 5 days of 4 μg/ml doxycycline treatment for induction of Cas9 then subclones isolated and screened for KO. *RNF168* was targeted with sequences TCGAAAAGGCGAGTTTATGC and ATCTGCATGGAAATCCTCG. Data is shown for a representative clone harboring sgRNA target sequence TCGAAAAGGCGAGTTTATGC. KO was confirmed by Sanger sequencing using primers; TGATACGCTTCTGGGCATAATA and ACCCCCTAACCTCTGAGAACTAT. MCF7 and MDA-MB-436 cells were transduced with lentivirus containing Cas9 and guide RNA in the lentiCRISPRv2 plasmid and selected using 2 μg/ml puromycin. The guide RNA targeting GCGACCATCCGGTTCCATGG was used to knockout *BARD1* in MCF7 cells and confirmed by Sanger sequencing using the following primers; GTGCCCTGCGAGTCCCTAT and CAAAAACTACCGTTTCAGTTGGAT. The guide RNA targeting sequence CCTCAACACGGACTCGGACA was used to knockout ABRAXAS in MCF7 cells and confirmed by sequencing with primers; CAGCAGAAGCGAAGGAGGA and GAGGGCTAATGCTGGAGAAGA. The guide RNA targeting GCCATCCAGTCCTCAAGGAG was used to knockout *53BP1* in MDA-MB-436 cells. The expression of 53BP1 in pooled population was confirmed by western blot using the nuclear extracts.

### BARD1, BRCA1, RNF168, PALB2 cloning and cell line generation

Full-length BARD1 (aa 1-777) was first cloned into the pENTR1A Gateway Entry vector (Thermo Fisher Scientific, A10462) with a GFP tag at the N-terminus. The RING domain (aa 34-126), ANK domain (aa 426-521), BRCT domain (aa 560-777), BUDR (R705A, D712A, Q715R) were deleted or mutated using the Q5 Site-Directed Mutagenesis Kit (New England Biolabs, E0554S). The GFP-BARD1 constructs were shuttled into a lentiviral destination vector containing a UBC promoter and a hygromycin resistance gene using the Gateway LR Clonase II Enzyme Mix (Thermo Fisher Scientific, 12538120). BRCA1-ΔRING (Δ1-127 aa, aka M128) and - ΔBRCT (Δ1496-1863 aa, aka 1495*) constructs were generated previously^23,29^. BRCA1-3A (I26A+L63A+K65A), -C61G, -ΔRING (Δ1-296 aa; M297), -ΔBRCT (Δ1559-1863 aa) and -ΔBRCT (Δ1815-1863 aa) constructs were generated using site directed mutagenesis or are described previously^23,29^. BARD1 aa 127-777 was linked using (Gly4Ser)2 to BRCA1 aa 128-1863 to generate a RING deficient BARD1/BRCA1 conjugated protein. The *RNF168* cDNA was a gift from Daniel Durocher and was cloned into the pENTR1A Gateway Entry vector with an N-terminal 3x FLAG tag. RNF168-ΔRING (aa 2-58) and -ΔPID (aa 524-571) domain deletions were generated by PCR and the R57D mutant by the QuickChange Lightning Site-Directed Mutagenesis Kit (Agilent, 210518) according to the manufacturer’s instructions. RNF168 constructs were shuttled into identical lentiviral destination vectors containing either a *UBC* or *CMV* promoter and a hygromycin resistance gene using the Gateway LR Clonase II Enzyme Mix. Full-length PALB2 cDNA was cloned into PENTR1A with an N-terminal HA tag, then shuttled into a lentiviral expression vector with a hygromycin resistance gene. BRCA1 RING domain (aa 2-104) was linked using (Gly4Ser)2 to the N-terminal of full-length PALB2 to generate the HA-RING-PALB2 construct. Lentivirus for each construct was produced in HEK293T cells (Takara Bio, 632180) using psPAX2 and VSV-G plasmids. Cell lines were infected with lentivirus using media with 10 μg/ml polybrene (Boston Bioproducts, BM-862W). Cells were then selected using the appropriate antibiotic (200 μg/ml hygromycin, 2 μg/ml puromycin, or 4 μg/ml blasticidin) or sorted for GFP expression. Construct expression was confirmed by Western blotting.

### RNAi experiments

Reverse transfection with Lipofectamine RNAimax (Thermo Fisher Scientific, 13778075) was performed according to the manufacturer’s instructions for siRNA experiments. The following siRNA were used: AllStars Negative Control (scrambled) siRNA (Qiagen, SI03650318), pooled *RAP80* targeted siRNA (Dharmacon J-006995-09-0002, J-006995-08-0002), pooled *RNF168* targeted siRNA (GACACUUUCUCCACAGAUAUU, CAGUCAGUUAAUAGAAGAAAUU, GUGGAACUGUGGACGAUAAUUU), pooled *RNF8* targeted siRNA (UGCGGAGUAUGAAUAUGAAUU, GGACAAUUAUGGACAACAAUU, AGAAUGAGCUCCAAUGUAUUU, CAGAGAAGCUUACAGAUGUUU), pooled *ABRAXAS* targeted siRNA (UAUUAGUGGUAACGUGAUA, CAGGGUACCUUUAGUGGUU).

### PARPi colony assays

Cells were seeded in 6 well plates into media containing DMSO or 1-10000 nM of rucaparib. Cells were fixed with 3:1 methanol:acetic acid after 2 weeks, stained with crystal violet, and colonies counted. IC50 calculations were performed with GraphPad Prism software using rucaparib response curves resulting from 3 or more independent experiments. In PARPi resistance assays, cells were seeded at varying densities into DMSO or 50 nM rucaparib and data presented as rucaparib treated colonies normalized to the DMSO treated control colonies for wells where distinct colonies could be counted. Rucaparib was provided by Clovis Oncology.

### Immunofluorescence microscopy and foci analyses

Cells were subject 10 Gy γ-irradiation (IR) then pre-extracted and fixed at 8 h post-IR. Pre-extraction was performed on ice for 5 min with cold cytoskeleton buffer (10 mmol/L PIPES pH 6.8, 100 mmol/L NaCl, 300 mmol/L sucrose, 3 mmol/L MgCl2, 1 mmol/L EGTA, 0.5% Triton X-100) followed by 5 min with cytoskeleton stripping buffer (10 mmol/L Tris-HCl pH 7.4, 10 mmol/L NaCl, 3 mmol/L MgCl2, 1% Tween 20 (v/v), 0.5% sodium deoxycholate). Pre-extracted cells were then fixed at room temperature for 10 min with 4% paraformaldehyde, washed with PBS, and treated for 10 min with 1% Triton X-100 in PBS. Primary antibodies were incubated overnight at 4 degrees in 5% goat serum in PBS. The following primary antibodies were used: Brca1 (gift from A. Nussenzweig), BRCA1 (Millipore, 07-434), RAD51 (Genetex, GTX100469), RAD51 (Abcam, ab133534), HA (Covance, MMS-101R), BARD1 (Santa Cruz Biotechnology, sc-74559), 53BP1 (Novus Biologicals, NB100), 53BP1 (Millipore, MAB3802), RPA32 (Cell Signaling, 2208), RPA32 (Sigma-Aldrich, NA18), RAP80 (Bethyl Laboratories, A300-763A), FLAG (Sigma-Aldrich, F1804), FLAG (Cell Signaling, 14793), CtIP (Millipore, MABE1060). To quantify PALB2 foci, cell lines expressing HA-PALB2 were generated and irradiated cells were stained using the anti-HA antibody. Experiments examining geminin (Abnova, H00051053-M01) staining did not use pre-extraction method (to preserve geminin staining). Here, the same protocol was used as above minus the pre-extraction step and beginning at the 4% paraformaldehyde fixation. Alexa Fluor or FITC conjugated secondary antibodies (Jackson ImmunoResearch Labs or Thermo Fisher Scientific) were incubated for 1h at room temperature and slides mounted using Vectashield antifade mounting media with DAPI (Vector Laboratories). Z-stack images were captured using a Nikon NIU Upright Fluorescence microscope and projection images generated using Nikon NIS Elements software. An ImageJ macro was used for quantification of percentage foci positive cells, foci size, and foci per nucleus. Nuclei were identified from the DAPI channel image using the analyze particles function of ImageJ after thresholding the image with the ImageJ median and subtract background functions applied. The identified nuclei were stored as ROI in the ImageJ ROI Manager to allow for characterization of foci within individual nuclei (ROI). Foci identification for each nucleus utilized the ImageJ subtract background, contrast, and gaussian blur functions followed by thresholding and the analyze particles function to generate a foci mask image. Threshold values were manually confirmed and applied consistently within each experiment. Foci per nucleus and foci size quantifications were direct outputs of the analyze particles function (particle count, particle average size) applied to the ROI of each nucleus on the foci mask images and are reported as counts and μm^2^, respectively. Foci positivity was defined as nuclei containing 5 or more RAD51 foci or 10 or more foci for other proteins as indicated in each figure. Percentage foci positive cells are presented as mean and S.E.M. from 3 independent experiments with the average of each biological replicate shown by open circle data points. Foci per nucleus and foci size data points represent the count and average foci size for individual foci positive nuclei across 3 independent experiments with the median values indicated. A minimum of 4 images and 100 nuclei were collected and analyzed per sample in each replicate. For geminin co-staining experiments, the ImageJ macro for foci analysis was adapted to include a step for thresholding of the geminin fluorescence image and geminin positivity assessed for individual nuclei.

### Western blotting and immunoprecipitation

Nuclear extracts were obtained from pelleted cells with the NE-PER Nuclear and Cytoplasmic Extraction Kit (Thermo Fisher Scientific, 78833) following instructions from the manufacturer and using protease and phosphatase inhibitors (Millipore, 524624, 524625, and 539131). Lysates were separated by SDS-PAGE, transferred to a PVDF membrane, and blocked in 5% nonfat milk in PBST. The primary antibodies were incubated overnight at 4 degrees in 5% nonfat milk in PBST. The following primary antibodies were used: HA (Cell Signaling, 2367), RNF168 (R&D Systems, AF7217), RNF168 (Millipore, ABE367), BRCA1 (Millipore, OP92), BARD1 (Bethyl Laboratories, A300-263A), GFP (Santa Cruz Biotechnologies, sc-9996), FLAG (Sigma-Aldrich, F1804), V5 (Bethyl Laboratories, A190-120A), PALB2 (Bethyl Laboratories, A301-246A), BRCA2 (Bethyl Laboratories, A303-434A), RAD51 (Santa Cruz Biotechnology, sc-8349), CtIP (Bethyl Laboratories, A300-488A), RAP80 (Bethyl Laboratories, A300-763A), ABRAXAS (Bethyl Laboratories, A302-180A), 53BP1(Millipore, MAB3802), RNF8 (Santa Cruz Biotechnology, sc-271462), Tubulin (Cell Signaling, 2148). HRP conjugated secondary antibodies (Millipore or R&D Systems) were incubated for 1h in 5% nonfat milk in PBST. V5 tag (Bethyl Laboratories, A190-120A) antibody was used for immunoprecipitation of BRCA1 complexes from 2 mg of nuclear extract using Pierce Classic IP Kit (Thermo Fisher Scientific, 26146) according to manufacturer’s instructions.

### Cell cycle analyses and RT-PCR

Exponentially growing cells were harvested and fixed with 50% ethanol at 4 degrees. Cells were washed with PBS and incubated in FxCycle PI/RNase staining solution (Thermo Fisher Scientific, F10797) for 30 min. Data were acquired using a BD FACScan flow cytometer and analyzed using FlowJo software. Total RNA was isolated from cell lines using the RNeasy Plus Mini Kit (Qiagen, 74134). The following primers are located within the 3x FLAG tag sequence and were used to quantify ectopic *FLAG-RNF168* expression: TACAAAGACCATGACGGTGATTA and AGTCTCGCTGCCTGAGATA. Quantification of ectopic of BRCA1 FL and -ΔBRCT expression was performed using primers located on the HA tag and N terminal of the *BRCA1* cDNA: CCTACGACGTGCCCGACTA and ATGGGACACTCTAAGATTTTCTGCA. Expression was normalized to *36B4* mRNA as previously described using q(quantitative) RT-PCR^23^.

### Statistical analyses

Statistical tests, significant p values and number of replicates are indicated in the figure legends. Nonparametric Mann-Whitney and unpaired, two-tailed t-tests were performed using GraphPad Prism software. R was used for chi-square goodness of fit tests comparing expected and observed genotypes.

## Acknowledgments

This work was supported by US National Institutes of Health (NIH) Grants R01CA214799, R01HL150190, and R01GM135293. J.J.K. was supported by an American Cancer Society - Tri State CEOs Against Cancer Postdoctoral Fellowship, PF-19-097–01–DMC, Ovarian Cancer Research Alliance and Phil and Judy Messing grant 597484, and T32 CA009035. We are grateful to Sean Hua and the FCCC Transgenic Mouse facility for help generating mice as well as Genomics, Cell Culture and Cell Sorting facilities. We thank Dr. Ross Chapman for helpful discussions, Dr. Kathy Cai and the FCCC histopathology service for pathological analyses of mouse organs, and Dr. Beth Handorf for guidance with statistical analyses.

## Author contributions

J.J.K., Y.W., P.P., J.B., A.J.B., designed and performed experiments J.J.K., Y.W., A.J.B., and N.J. analyzed and interpreted data. J.J.K., Y.W., and N.J. wrote the manuscript. N.J. supervised the project.

---

**Figure S1.**
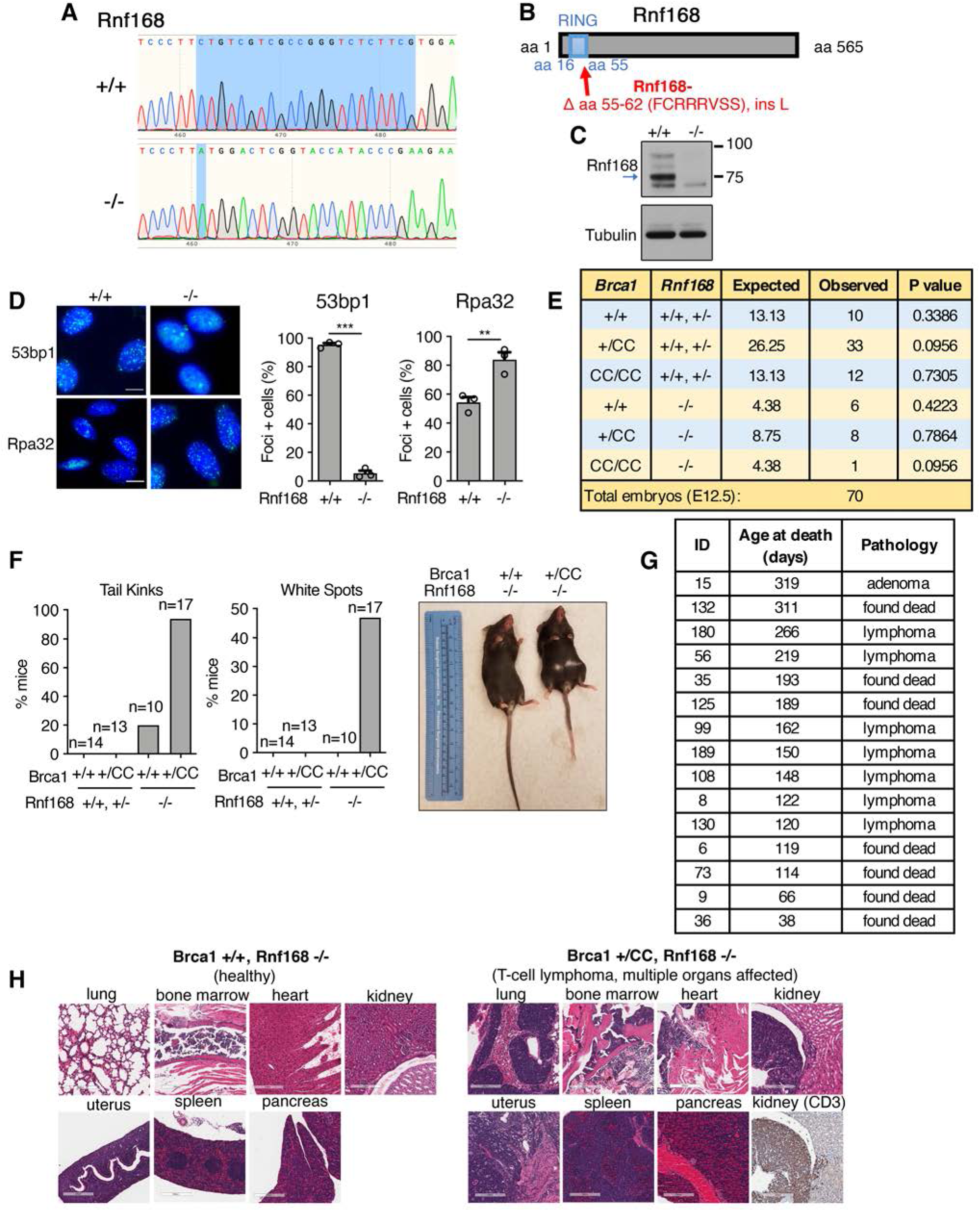
Phenotypes associated with *Rnf168*^-^ and *Brca1^CC^* mice, related to Fig. 1. **(A)** Electropherogram showing *Rnf168* wild-type (+/+) and mutant (-/-) alleles. The *Rnf168^-^* allele has a deletion of CTGTCGTCGCCGGGTCTCTTCG and insertion A at c.165. **(B)** Cartoon showing the impact of the above *Rnf168^-^* mutation on the Rnf168 protein. *Rnf168^-^* generates an in-frame deletion of amino acids (aa) 55-62 plus an insertion of an L. **(C)** Rnf168 protein expression was assessed in *Rnf168^+/+^* and *Rnf168^-/-^* MEFs using an antibody recognizing an epitope spanning aa 423-565. Despite the *Rnf168^-^* allele generating an in-frame deletion no protein was detected by Western blotting, likely from protein destabilizing effects of the deletion mutation that is proximal to the Ring domain. **(D)** 53bp1 and Rpa32 IRIF were measured in *Rnf168^+/+^* and *Rnf168^-/-^* derived MEFs. *Left*; representative images (scale bar, 10 μm). *Right*; mean and S.E M. percentage (≥10) foci positive cells, n=3 biological replicates. *** p < 0.001, ** p < 0.01, (unpaired, two-tailed t-tests). **(E)** Live embryos from *Brca1^+/CC^, Rnf168^+/-^ x Brca1^+/CC^, Rnf168^+/-^* crosses were collected and genotyped at E12.5. The numbers of expected and observed live embryos is shown. P value are obtained from chi-square goodness of fit tests for the binomial of each genotype. **(F)** Mice with the indicated genotypes were recorded for the number born with tail kinks and white spots and expressed as a percentage of the number of mice evaluated for each of the indicated genotypes. *Right*; photograph of representative mice. **(G)** Table describing age and available pathology at time of death associated with *Brca1^+/CC^, Rnf168^-/-^* mice. **(H)** The indicated organs and tissues were H&E stained and subject to pathological inspection. Images are shown from a representative healthy *Brca1^+/+^, Rnf168^-/-^* mouse and *Brca1^+/CC^, Rnf168^-/-^* mouse with lymphoma detected in multiple organs (scale bars, 200 μm). CD3 immunohistochemical staining of the lymphocytic infiltration of the kidney (bottom right) indicates T-cell lymphoma.

**Figure S2.**
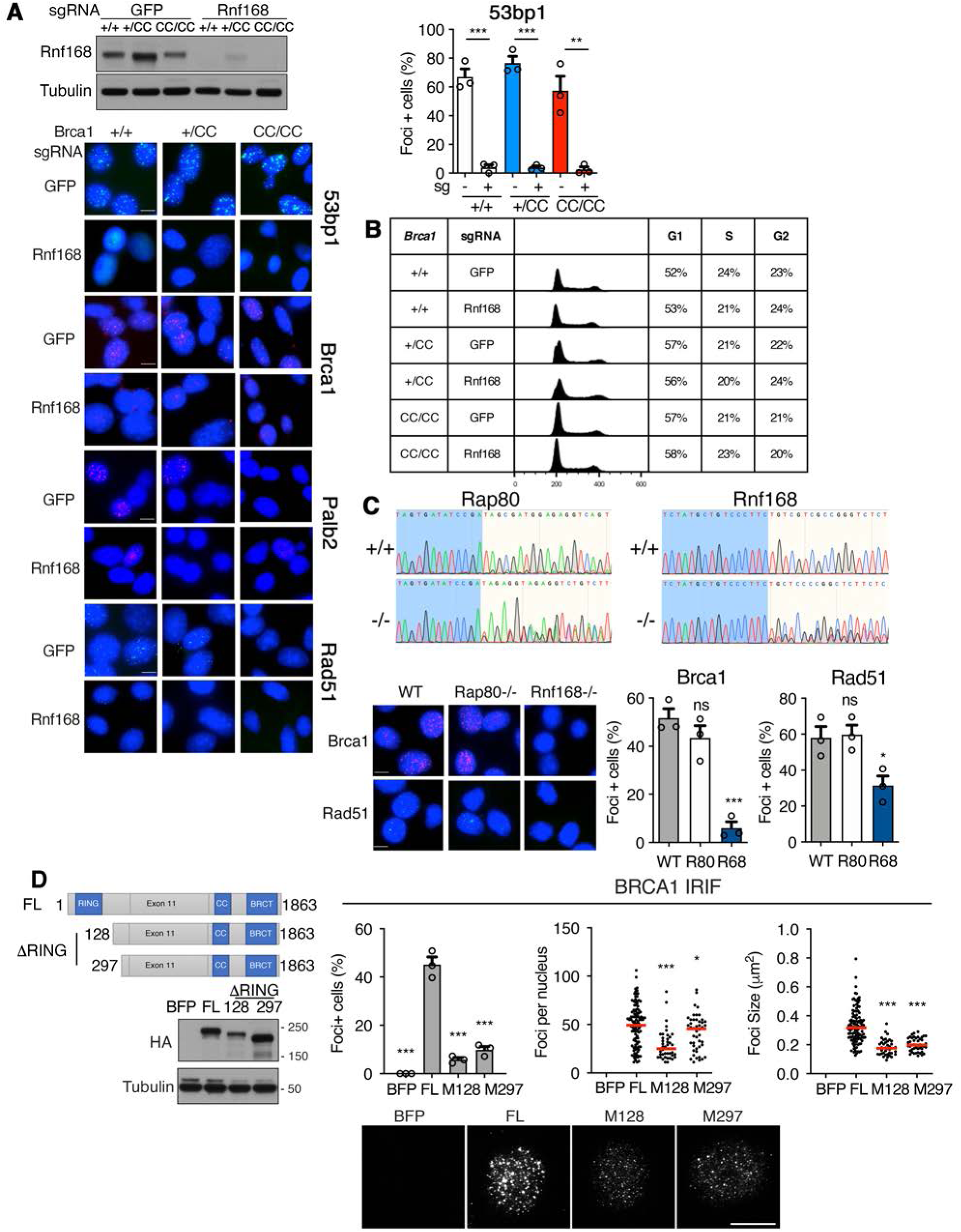
Characterization of *Rnf168*^-^ and *Brca1^CC^* MEFs, related to Fig. 1&2. **(A)** *Brca1^+/+^*, *Brca1^+/CC^*, and *Brca1^CC/CC^* MEFs were incubated with lentivirus expressing sgRNA targeting *GFP* (control) or *Rnf168* and cells collected for Western blotting to confirm loss of Rnf168 expression. *Below*; representative pictures of cells from Fig. 1F (scale bar, 10 μm). 53bp1 IRIF were quantified as described in Fig. 1F. **(B)** To confirm that differences in foci formation were not caused by significant changes in cell cycle distribution, cells from A were assessed for cell cycle fractions using propidium iodide (PI) staining and flow cytometry. **(C)** MEFs were subject to sgRNA targeting *Rnf168, Rap80* or *GFP* (WT), clones established with frameshifting mutations and assessed for Brca1 and Rad51 IRIF. *Above*; Electropherograms and frameshift mutations are shown for *Rap80^-/-^* and *Rnf168*^-/-^ clones. *Below*; representative pictures (scale bar, 10 μm) and mean and S.E.M. percentage BRCA1 (≥10) foci and RAD51 (≥5) foci positive cells. MEFs n=3 biological replicates. *** p < 0.001, * p < 0.05, ^ns^ not significant compared to WT MEFs (unpaired, two-tailed t-tests). **(D)** *Left*; cartoon showing BRCA1 protein domains and truncations with aa numbers indicated. *Below*; HA Western blot of MDA-MB-436 cells expressing BFP control, HA-BRCA1-full-length (FL), HA-BRCA1-ΔRING with aa start at M128 and M297. *Right*; mean and S.E.M. percentage foci positive cells; foci positive cells were assessed for the number of foci present in a single nucleus, red line: median value; foci positive cells were also assessed for the mean size of foci present per nuclei, n=3 biological replicates. Foci positive cells were determined to be those with ≥10 foci/nuclei. *** p < 0.001, ** p < 0.01, * p < 0.05, ^ns^ not significant compared to FL (percentage positive cells: unpaired, two-tailed t-tests size; number and size per nucleus: nonparametric Mann-Whitney tests). *Below*; representative images (scale bar, 10 μm).

**Figure S3.**
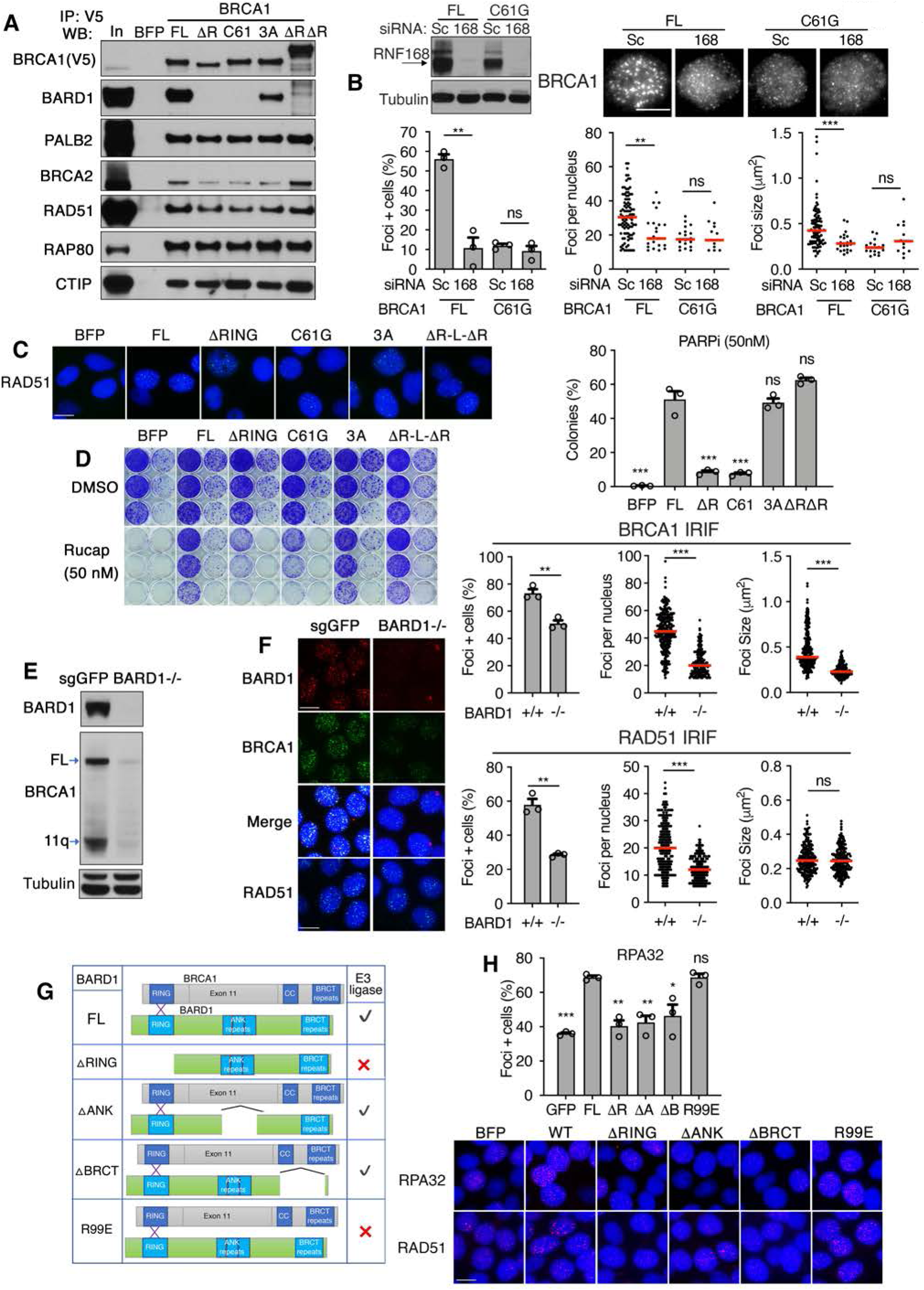
Assessment of BRCA1 and BARD1 foci formation, related to Fig. 3. **(A)** Cartoon showing BRCA1-full-length (FL), BRCA1-ΔRING (ΔR), BRCA1-C61G (C61), BRCA1-I26A+L63A+K65A (3A), BRCA1-ΔRING-L-BARD1-ΔRING (ΔRΔR) predicted effects on BARD1 interaction and E3 ligase activity. *Right*; ectopic V5-tagged constructs expressed in MDA-MB-436 cells were immunoprecipitated and subject to Western blotting for the indicated proteins. **(B)** MDA-MB-436 cells expressing BRCA1-FL or BRCA1-C61G were subject to scrambled (Sc) and RNF168-targeting siRNA followed by IR and HA foci formation measured by immunofluorescence. *Left*; Western blot showing RNF168 depletion. *Above*; representative cells (scale bar, 10 μm). *Below*; foci analyses as described in Fig. 2B, n=3 biological replicates. **(C)** Representative RAD51 foci images from Fig. 3C (scale bar, 10 μm). **(D)** Representative 6-well plates showing colony formation of increasing cell densities that were seeded in the presence of either DMSO or 50 nM rucaparib. Mean and S.E.M. colonies that grew in the presence of rucaparib calculated as a percentage of those that grew in the presence of DMSO, n=3 biological replicates. **(E)** MCF7 cells were subject to sgRNA targeting *GFP* or *BARD1*, individual colonies generated and *BARD1^-/-^* cells identified by Sanger sequencing. The effects of *BARD1* KO on BARD1, BRCA1 full-length and BRCA1-Δ11q protein expression were assessed by Western blotting. **(F)** MCF7 cells harboring sgRNA targeting *GFP* and a *BARD1^-/-^* clone were assessed for BARD1, BRCA1 and RAD51 IRIF. *Left*; representative pictures (scale bar, 10 μm). *Right;* BRCA1 and RAD51 IRIF. *Left*; mean and S.E.M. percentage foci positive cells; *middle*, foci positive cells were assessed for the number of foci present in a single nucleus, red line: median value; *right*, foci positive cells were assessed for the mean size of foci present per nuclei, n=3 biological replicates. Foci positive cells were determined to be those with ≥10 foci/nuclei for BRCA1 and ≥5 foci/nuclei for RAD51. *** p < 0.001, ** p < 0.01, * p < 0.05, ^ns^ not significant (percentage positive cells: unpaired, two-tailed t-tests size; number and size per nucleus: nonparametric Mann-Whitney tests). **(G)** Cartoon showing BARD1-full-length (FL), BARD1-ΔRING, BARD1-ΔANK, BARD1-ΔBRCT, and BARD1-R99E predicted effects on BARD1 interaction and E3 ligase activity used in Fig. 3E-H. **(H)** Representative pictures as well as RPA32 IRIF from cells treated as in Fig. 3G (scale bar, 10 μm).

**Figure S4.**
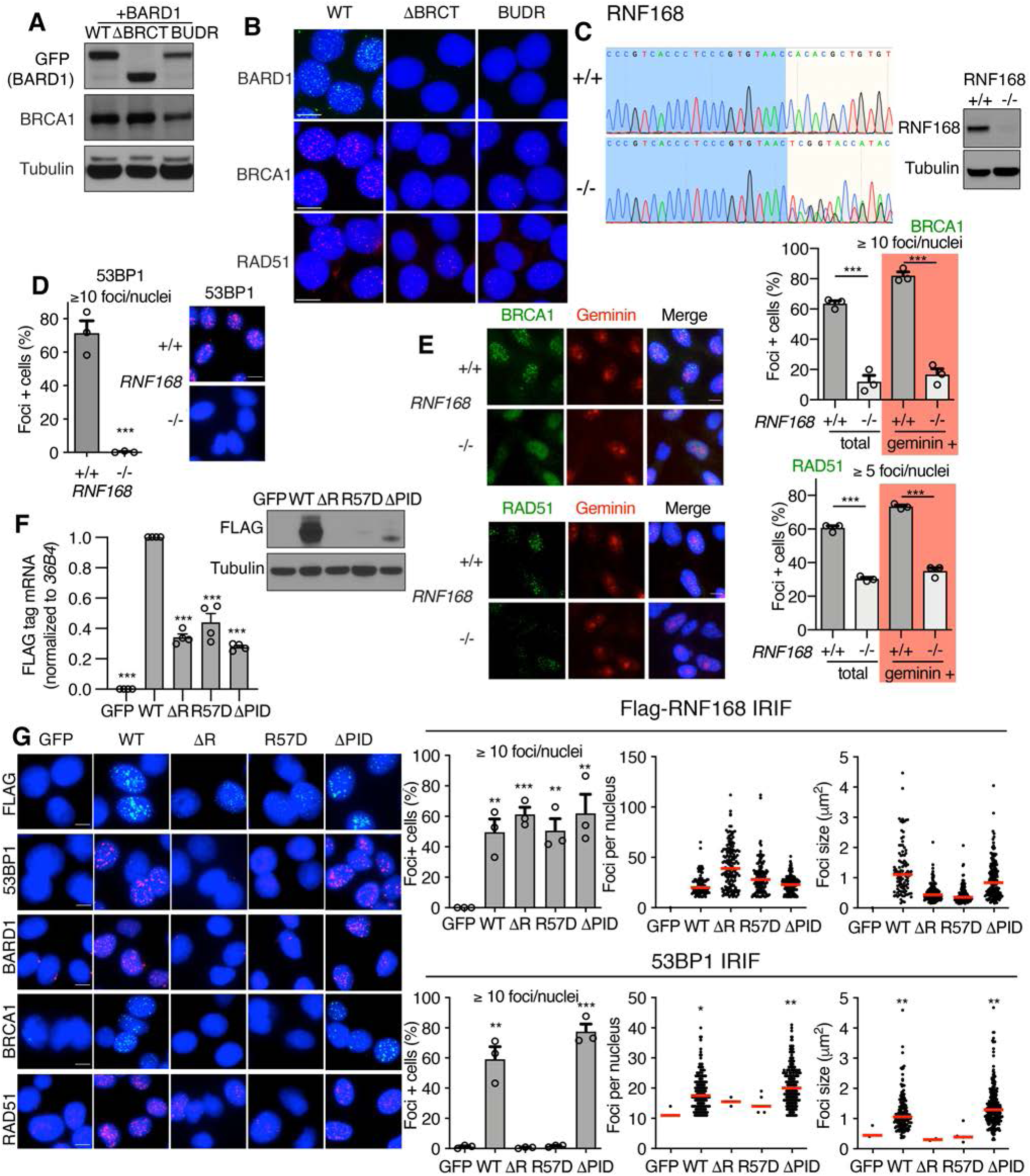
Characterization of *RNF168* KO cells, related to Fig. 4. **(A)** MCF7 *BARD1^-/-^* cells expressing GFP-BARD1-WT, GFP-BARD1-ΔBRCT, or GFP-BARD1-BUDR (R705A+D712A+Q715R) were assessed for GFP and BRCA1 expression by Western blotting. **(B)** Representative foci images from Fig. 4A and B (scale bars, 10 μm). **(C)** Electropherograms of *RNF168* DNA sequences from MDA-MB-231 WT and *RNF168^-/-^* cell lines are shown. The *RNF168^-/-^* cell line contains two alleles with frameshifting deletions. *Right*; Western blot showing loss of RNF168 expression. **(D)** Cells from C were assessed for 53BP1 IRIF. Representative images of cells are shown. Mean and S.E.M. percentage foci positive cells are shown from n=3 biological replicates. *** p < 0.001 (unpaired, two-tailed t-tests). **(E)** Cells from C were assessed for BRCA1 and geminin or RAD51 and geminin staining in the absence of pre-extraction by immunofluorescence. Representative images are shown. Mean and S.E.M. percentage foci positive cells as well as foci positive cells that were also geminin positive are shown from n=3 biological replicates. Geminin indicates S/G2 cell cycle fractions. *** p < 0.001, ** p < 0.01, * p < 0.05, ^ns^ not significant (unpaired, two-tailed t-tests). **(F)** RNF168 ectopic UBC promoter constructs were assessed for Flag mRNA and protein expression by qRT-PCR (*left*) and Western blotting (*right*), respectively. **(G)** Representative images from Fig. 4E as well as Flag-RNF168 and 53BP1 IRIF analyses as described in Fig. 4E. *** p < 0.001, ** p < 0.01, * p < 0.05, ^ns^ not significant compared to GFP expressing cells for all panels (unpaired, two-tailed t-tests). Scale bars, 10 μm for all panels.

**Figure S5.**
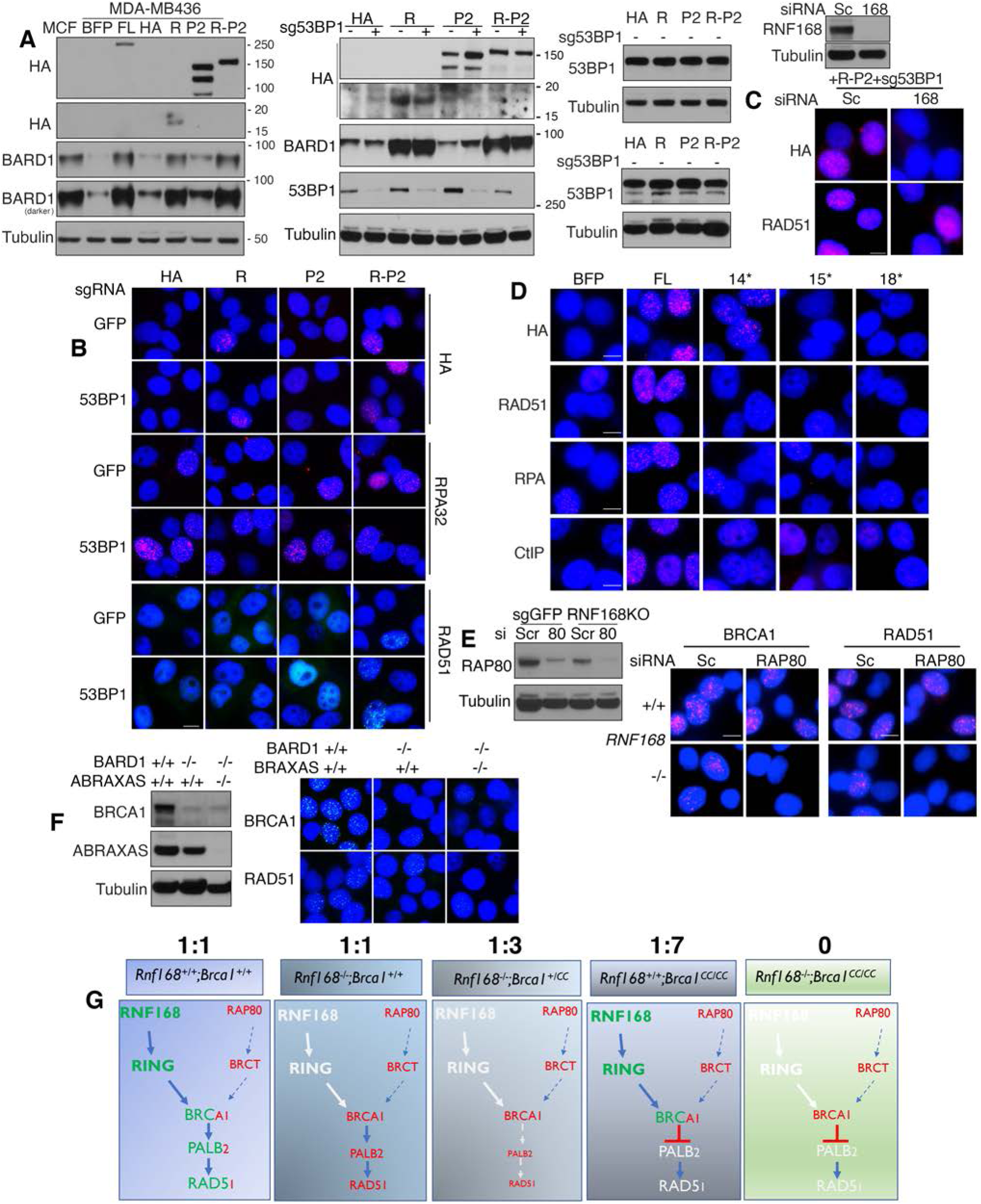
RING constructs and the BRCA1-A complex, related to Fig. 4&5. **(A)** Associated with Fig. 4G: *Left*, MDA-MB-436 cells expressing HA-empty vector (HA), HA-BRCA1 RING domain (R), HA-PALB2 (P2), or -HA-BRCA1 RING domain-PALB2 (R-P2) were subject to sgRNA targeting GFP or 53BP1 and assessed for the indicated protein expression by Western botting. To determine the relative effects of RING constructs on BARD1 expression, protein levels were compared to MCF7 as well as MDA-MB-436 expressing ectopic BFP and full-length BRCA1. *Middle*, Western blots showing the effects of sg53BP1 on 53BP1 protein expression in the indicated cell lines. *Right*, Two sets of additional cell lysates were generated for 53BP1 wild-type cells to assess whether 53BP1 protein levels were similar between cell lines. **(B)** Representative images from Fig. 4G. **(C)** Representative images and western blot associated with Fig. 4I. **(D)** Representative images associated with Fig. 5B. **(E)** Representative images and western blot associated with Fig. 5E. **(F)** Representative images and western blot associated with Fig. 5F. **(G)** Model for impact of *Brca1^CC^* and *Rnf168^-^* alleles and the supporting role of Rap80-Abraxas on Palb2 and Rad51 loading and development in mice. The Mendelian birth ratios of live mice are indicated above.

